# Urea nitrogen recycling via gut symbionts increases in hibernators over the winter fast

**DOI:** 10.1101/2021.02.24.432731

**Authors:** Matthew D. Regan, Edna Chiang, Yunxi Liu, Marco Tonelli, Kristen M. Verdoorn, Sadie R. Gugel, Garret Suen, Hannah V. Carey, Fariba M. Assadi-Porter

**Author notes:** Département de Sciences Biologique, Université de Montréal, Montréal, QC, H2B 0B3.

## Abstract

Hibernation is a mammalian strategy that uses metabolic plasticity to reduce energy demands and enable long-term fasting. Fasting mitigates winter food scarcity but eliminates dietary nitrogen, jeopardizing body protein balance. Here, we reveal gut microbiome-mediated urea nitrogen recycling in hibernating 13-lined ground squirrels (TLGS). Ureolytic gut microbes incorporate urea nitrogen into organic compounds that are absorbed by the host, with the nitrogen reincorporated into the TLGS protein pool. Urea nitrogen recycling is greatest after prolonged fasting in late winter, when urea transporter abundance in gut tissue and urease gene abundance in the microbiome are highest. These results reveal a functional role for the gut microbiome in hibernation and suggest mechanisms by which urea nitrogen recycling contributes to protein balance in other monogastric animals, including humans.

**One Sentence Summary:** Ground squirrels and their gut symbionts benefit from urea nitrogen recycling throughout hibernation.

## Main Text

Hibernation is a complex trait employed by animals to survive seasonal food scarcity. The hallmark of hibernation is torpor, a metabolically depressed state in which the hibernator’s rate of metabolic fuel use is reduced by up to 98% relative to active season rates (*1*). Torpor enables seasonal hibernators like the 13-lined ground squirrel (*Ictidomys tridecemlineatus*; TLGS) to fast for the ~6-month winter hibernation season, effectively solving the problem of winter food scarcity; however, this fasting deprives the TLGS of a dietary nitrogen source, which may jeopardize protein balance.

Despite the lack of dietary nitrogen and prolonged inactivity, hibernators lose little muscle mass and function during winter (*2*). Moreover, late in hibernation, TLGS elevate muscle protein synthesis rates to active season levels (*3*). It is unknown how hibernators preserve tissue protein during hibernation, but one hypothesis suggests that they harness the ureolytic activity of gut microbes to recycle urea nitrogen back into their protein pools through urea nitrogen salvage (UNS; Fig. 1) (*4*). UNS is present in ruminant and non-ruminant animals (*5*), but little direct evidence exists for its use by mammalian hibernators (*6*).

**Fig. 1.**
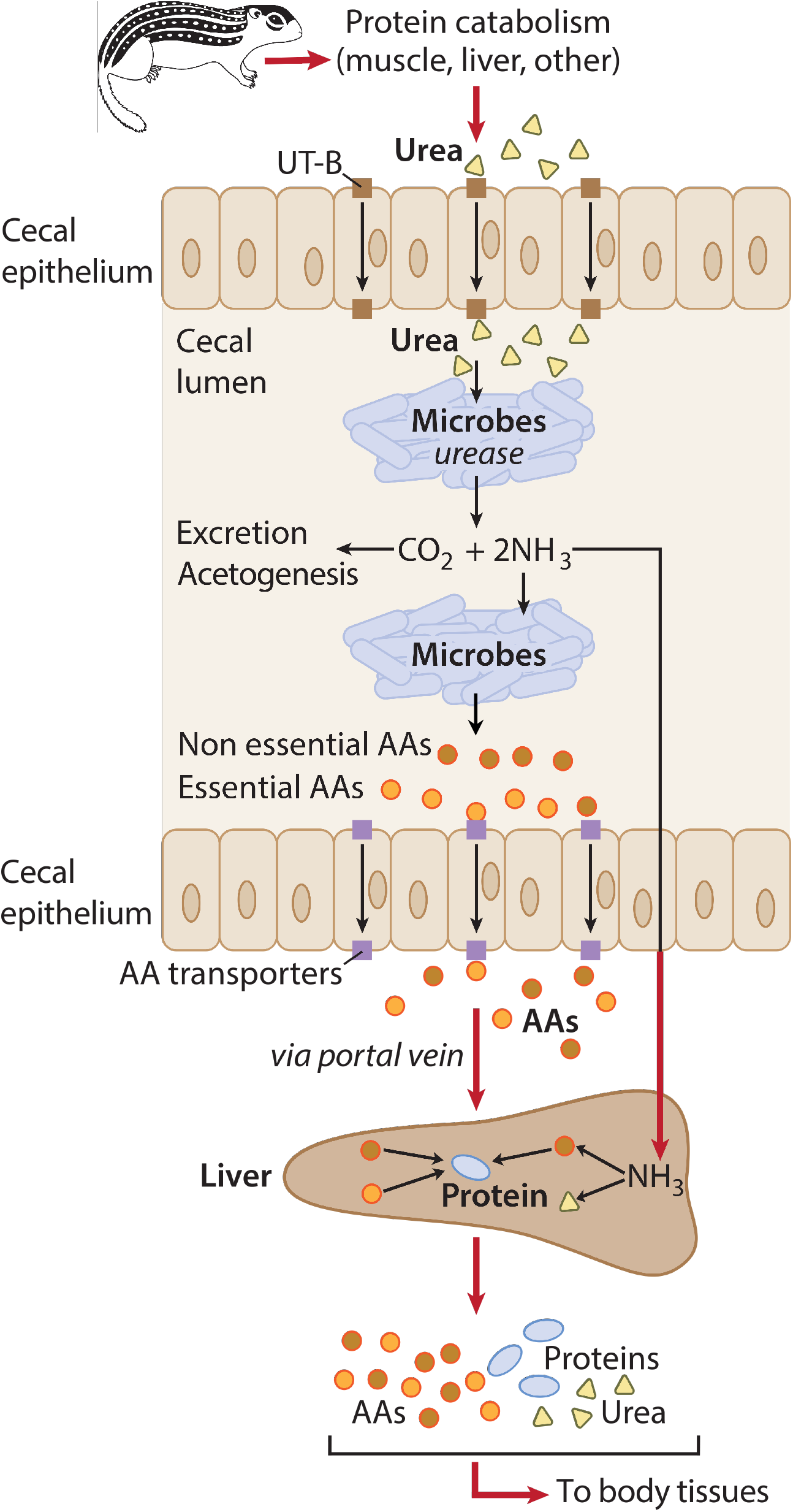
Urea nitrogen salvage (UNS) mechanism. Urea (yellow triangles), endogenously produced *via* protein catabolism, is transported *via* epithelial urea transporters (UT-B; brown squares) from blood into cecal lumen where, in the presence of urease-expressing gut microbes, it is hydrolyzed into CO_2_ and ammonia (NH_3_). CO_2_ can be excreted by the host and/or fixed by microbes. Microbes use NH_3_ to synthesize microbial amino acids (AA; circles), which are incorporated into their proteome or absorbed by the host *via* ceco-colonic AA transporters (purple squares) (*21*). Alternatively, NH_3_ is absorbed by the host and converted into AAs and/or urea in the liver. Ultimately, the AAs are used to synthesize protein (blue ovals) in host tissues, recycling the urea nitrogen.

We hypothesized that during hibernation, urea nitrogen is recycled into TLGS protein pools *via* UNS. We tested this hypothesis using three seasonal TLGS groups: Summer (active), Early Winter (one month into hibernation/fasting), and Late Winter (3-4 months into hibernation/fasting). Early and Late Winter TLGS were observed during induced interbout arousals, when metabolic rates and body temperatures were at euthermic levels. Each seasonal group contained TLGS with intact and antibiotic-depleted gut microbiomes. For each group, we administered two intraperitoneal (i.p.) injections of ^13^C,^15^N-urea (~7 d apart depending on season; unlabeled urea used as control; Table S2) and then examined the critical steps of UNS (Fig. 1).

UNS begins with hepatic urea synthesis and transport into the blood. Urea that is not excreted by the kidneys can be transported into the gut lumen *via* epithelial urea transporters (UT-B; (*7*)) where, in the presence of urease-expressing microbes, it is hydrolyzed into ammonia and CO_2_. We measured lower plasma urea concentrations in Early and Late Winter than Summer TLGS (Fig. 2A; P=0.0002), suggesting less urea is available for hydrolysis during hibernation, when UNS would be most beneficial. However, cecal UT-B abundance was ~3-fold higher in Late Winter than Summer TLGS (Fig. 2B; Season factor P=0.0002), suggesting that a greater capacity to transport urea into the gut lumen may partially compensate for (or contribute to) the lower plasma urea concentrations in winter. The mechanism underlying greater UT-B abundance in winter TLGS may involve luminal ammonia, which in other mammals inhibits epithelial UT-B expression (*8*). Commensurate with this, luminal ammonia levels were ~3-fold lower in hibernation than Summer ((*9*); Fig. S1; P<0.0001).

**Fig. 2.**
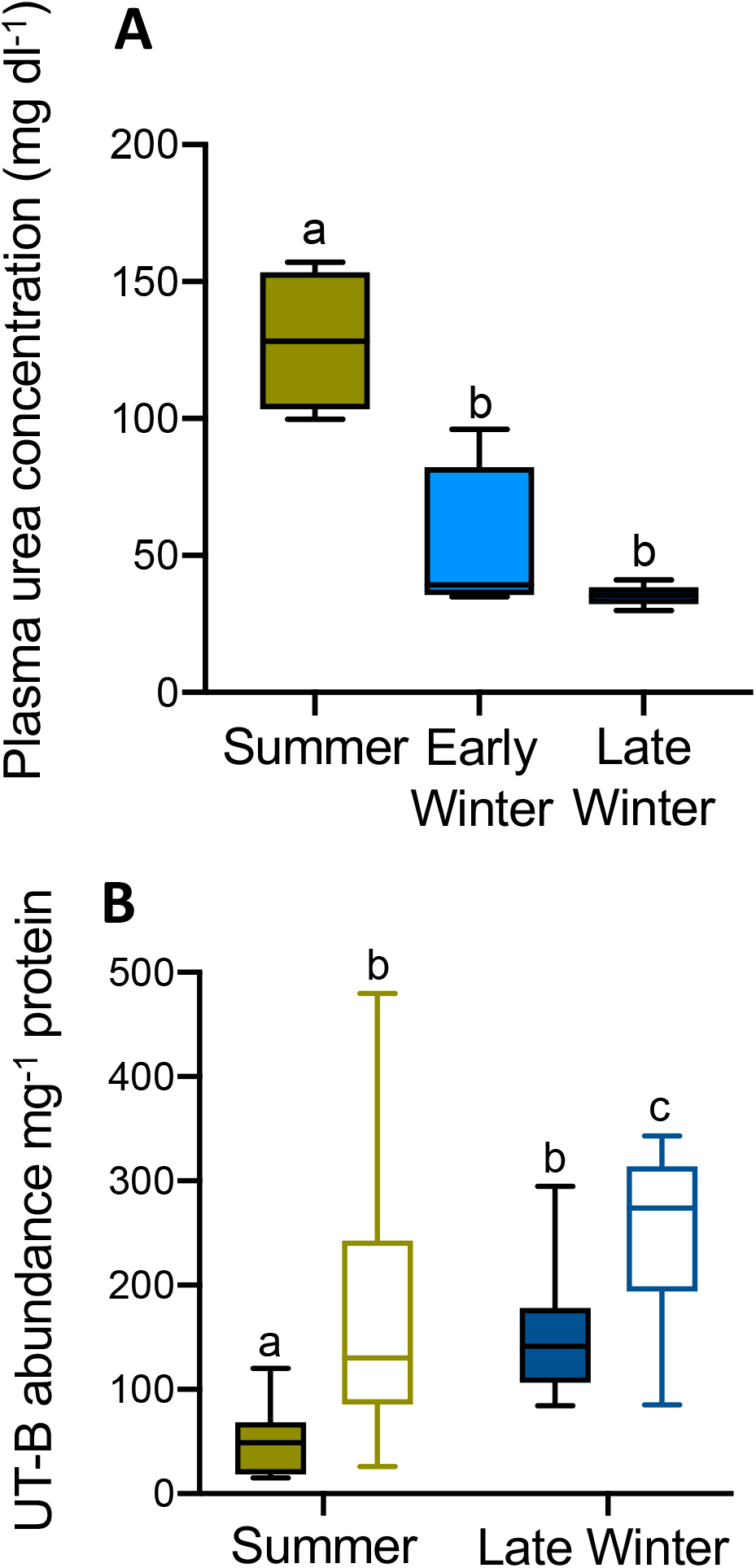
Plasma [urea] and cecal urea transporter (UT-B) expression. (A) Plasma [urea] in urea-treated TLGS (n=4-5). (B) UT-B protein abundance per mg protein in cecum tissue of non-urea-treated TLGS with intact (filled bars) and depleted (open bars) gut microbiomes (n=12). Dissimilar letters indicate significant differences (P<0.05).

Next, we measured microbial ureolytic activity *in vivo* using stable isotope breath analysis, where an increase in ^13^CO_2_:^12^CO_2_ (δ^13^C) following ^13^C,^15^N-urea injection indicates microbial ureolysis. Breath δ^13^C increased following ^13^C,^15^N-urea injection (Fig. 3A-C) in microbiome-intact, but not microbiome-depleted, TLGS (Fig. 3D; Microbiome factor P<0.0001), confirming microbial ureolytic activity. Ureolysis was greatest in Summer (Fig. 3D, Season factor P<0.0001; Fig. S1, P=0.028), consistent with the ~10-fold greater abundance of gut bacteria in summer *versus* hibernating ground squirrels (*10*) and the seasonal shift in microbial CO_2_ fixation (*e.g.*, microbial acetogenesis; Fig. S2). Nevertheless, the significantly elevated breath δ^13^C values in Early and Late Winter TLGS following urea injection indicate that ureolysis continues throughout hibernation.

**Fig. 3.**
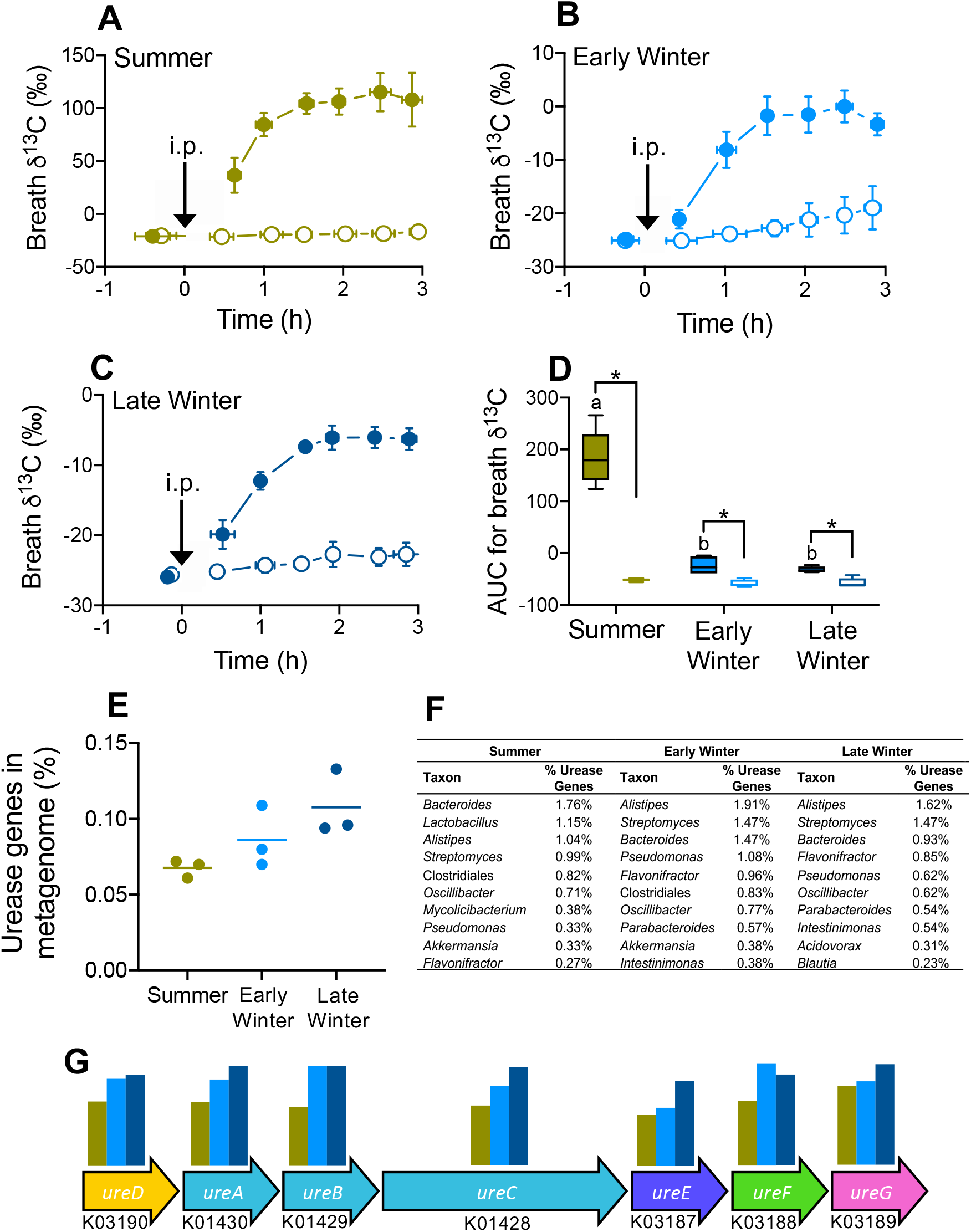
Gut microbial ureolysis and metagenome. Mean breath δ^13^C for (A) Summer, (B) Early Winter, and (C) Late Winter TLGS treated with ^13^C,^15^N-urea. X- and Y-error bars represent SEM and closed and open symbols represent TLGS with intact and depleted microbiomes, respectively. Arrow indicates ^13^C,^15^N-urea intraperitonial (i.p.) injection at time zero. (D) Mean area-under-the-curve (AUC) values for data in (A), (B), and (C), where asterisks indicate significant differences between microbiome groups within a season and dissimilar letters indicate significant differences between seasonal groups among microbiome-intact TLGS (n=5 A-D except n=4 for Early Winter microbiome-depleted). Seasonal effect on (E) the percentage of urease genes in the gut metagenome and (F) the top ten microbial taxa from taxonomic classification of urease genes. Urease operon (G) including each gene’s KEGG ontology ID and seasonal relative abundances.

Despite reduced bacterial abundance during hibernation (*10*), the metagenomes of hibernating TLGS trended towards a higher percentage of urease genes than those of Summer TLGS (Fig. 3E, P=0.083). This included six of the seven urease-related genes (Fig. 3G), suggesting that during hibernation, a higher percentage of microbes have the potential to hydrolyze urea. Indeed, of the ten microbial taxa with the greatest detectable genomic representation of urease genes in Late Winter (Fig. 3F), five are members of taxa that increase in relative abundance throughout hibernation (*9*).

To benefit the host, microbial ureolytic activity would need to provide urea-derived compounds such as amino acids (AAs), peptides and/or ammonia (the latter convertible to AAs in liver). Using 2-dimensional ^1^H-^15^N spectroscopy, we found that more ^15^N was incorporated into cecal content and liver metabolomes of microbiome-intact than -depleted TLGS (Fig. 4A,B), a trend that, with few exceptions, also held for specific compounds including ammonia, glutamine, alanine, lysine, and valine (Fig. 4A,B). ^15^N-metabolite levels were also affected by season. In cecum content, Early and Late Winter metabolite levels were generally lower than in Summer, whereas for the liver, Early and Late Winter metabolite levels were generally higher than in Summer and typically highest in Late Winter (Fig. 4A,B). In contrast, for muscle, there was little indication that metabolite ^15^N incorporation was affected by the presence of a microbiome. Given the significant ^15^N incorporation into muscle protein (see below), this may result from the timing of ^13^C,^15^N-urea dosing and tissue sampling protocols. By the time of tissue sampling, the ^15^N-AAs from the initial ^13^C,^15^N-urea dose may have been incorporated into muscle protein, thus explaining the ^15^N-protein results (Fig. 4C), while the 3 h between the second dose and tissue sampling may have been too brief for ^15^N-metabolites to appear in the muscle. The muscle ^15^N-metabolite results could therefore represent microbiome-independent background ^15^N levels, which is consistent with equivalent ^15^N-metabolite abundances in the muscles of TLGS treated with labeled and unlabeled urea, and with intact and depleted microbiomes (Table S2).

**Fig. 4.**
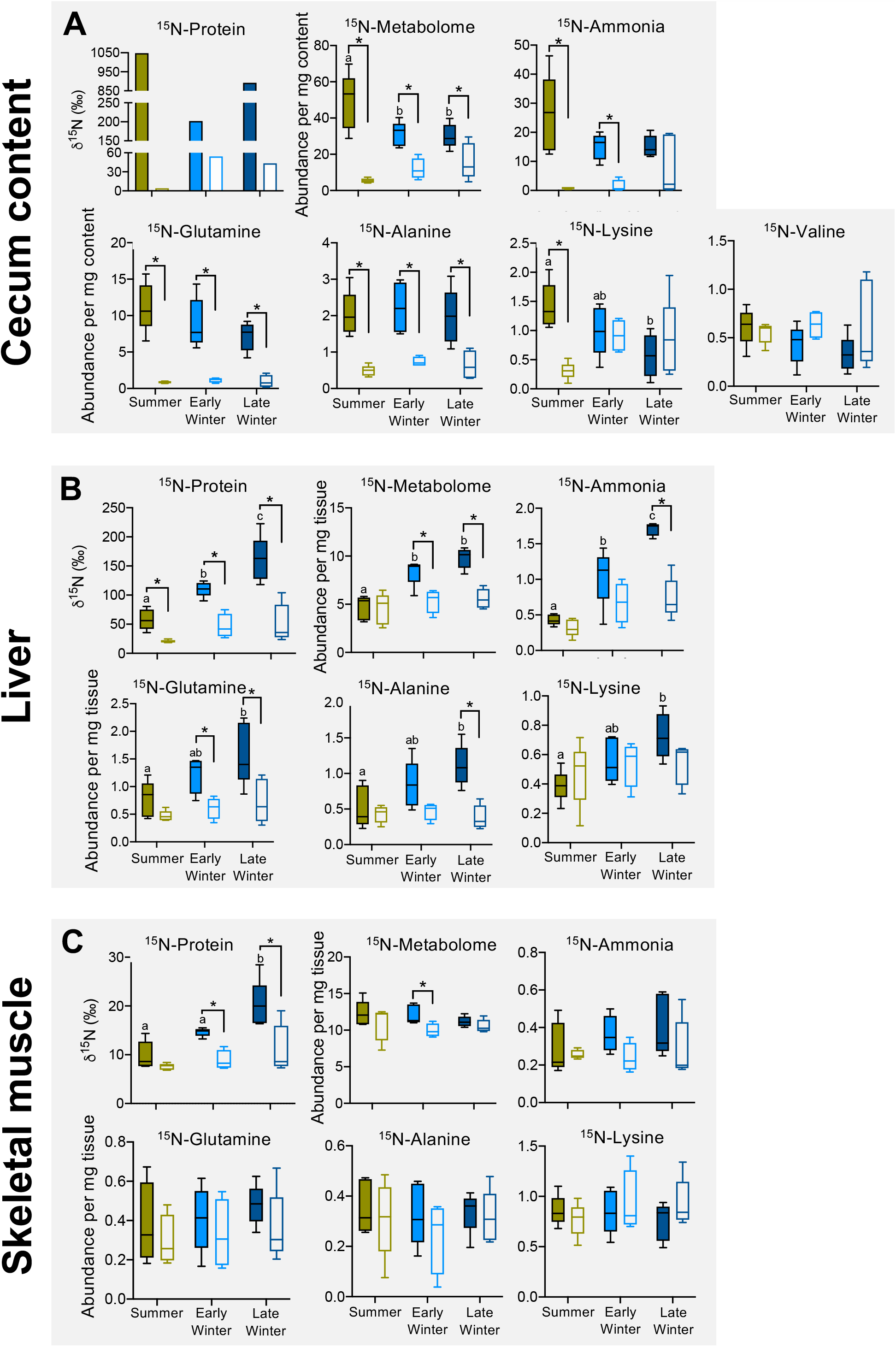
^15^N-urea nitrogen incorporation into metabolite and protein pools of gut microbiome and host tissues. (A) Cecal content, (B) liver, and (C) muscle (quadriceps) of microbiome-intact (filled bars) and -depleted (open bars) TLGS. For each panel, ^15^N-Protein portrays ^15^N-incorporation into protein content, ^15^N-Metabolome portrays ^15^N-incorporation into the total pool of compounds (identifiable and non-identifiable), and remaining plots portray ^15^N-incorporation into the compound named atop the plot. Metabolomic data are relative abundances in arbitrary units. Asterisks indicate significant difference between microbiome-intact and -depleted groups within a season, and dissimilar letters indicate significant seasonal difference between microbiome-intact groups. See Supplementary Material for statistical results details. n=5 for all data except n=4 for Early Winter microbiome-depleted and n=1 (pooled samples) for cecum content ^15^N-protein.

The ultimate benefit of UNS is urea nitrogen incorporation into host protein. Using isotope ratio mass spectrometry, we found that microbiome-intact TLGS incorporated significant ^15^N into liver and muscle protein relative to microbiome-depleted TLGS (Fig. 4B,C), thereby verifying that UNS is microbially dependent and benefits the host. Moreover, ^15^N incorporation into liver and muscle protein was 2-to 3-fold higher in Late Winter than Summer, demonstrating that UNS is most beneficial late in the winter fast. We also found large amounts of ^15^N incorporated into cecal content protein (indicative of microbial protein; Fig. 4A) in microbiome-intact TLGS, indicating nutritional benefits for the gut microbiome. However, unlike in liver and muscle, microbial protein ^15^N incorporation was highest in Summer (Fig. 4A), when microbial abundance is highest.

Interestingly, ^15^N incorporation into host protein was highest when ureolytic activity was lowest (i.e., Late Winter). This is explained by our observations at multiple UNS steps. First, urea transport capacity into the gut is likely increased during hibernation *via* elevated UT-B abundance (Fig. 2). Second, despite lower overall ureolytic activity (Fig. 3D), the winter microbiome is proportionally more ureolytic than the summer microbiome. This is evidenced by elevated urease gene percentages (Fig. 3E) and ~2-fold higher urease activity levels (Fig. S1B) than the breath δ^13^C changes (Fig. 3D) and 10-fold reduction in microbial abundance during winter (*10*) predict. Third, higher bacterial death in winter (*10*) could liberate relatively more bacterial contents for host absorption during hibernation, including microbial urea nitrogen-enriched metabolites (Fig. 4A). The higher mass-specific AA absorption rates of intestinal tissues from hibernating *versus* summer TLGS (*11*) may also contribute to the elevated appearance of ^15^N-AAs in liver (Fig. 4B).

Perhaps most crucially, hepatic conversion of urea-derived ammonia into glutamine *via* glutamine synthetase (GS) may be enhanced during hibernation. Numerous lines of evidence support this, including elevated liver ^15^N-ammonia and ^15^N-glutamine levels in Early and Late Winter (Fig. 4B), sustained liver GS activity during hibernation (Fig. S3), and reduced urea cycle enzymes and intermediates (*12*, *13*). This implies that, during hibernation, ammonia conversion to glutamine is favored over conversion to urea. Indeed, in marginally protein-deficient rats, hepatic ammonia processing shifts from urea toward glutamine production (*14*), and in fasting hibernating arctic ground squirrels, ammonia from muscle protein breakdown is directed away from the urea cycle towards AA formation (*13*). Despite this shift in ammonia processing, urea production continues during hibernation (albeit at lower rates than during summer), with plasma concentrations highest during interbout arousal and then falling to minimum levels upon torpor reentry (Fig. S4; (*15*)). Much of this urea is likely transferred to the gut *via* UT-B, where it becomes available for microbial ureolysis. Thus, nitrogen recycling in hibernators involves both endogenous (*13*) and microbe-dependent mechanisms, underscoring the importance of maintaining nitrogen balance during hibernation.

UNS provides two major benefits for hibernating animals. First, UNS augments protein synthesis during hibernation, when dietary nitrogen is absent. UNS appears especially important late in hibernation, just prior to TLGS’s emergence into its breeding season (*3*, *16*). By facilitating protein synthesis and subsequent tissue function, UNS may provide a selective advantage during mating and thereby influence biological fitness. Second, UNS may enhance water conservation during hibernation. UNS diverts urea away from the kidney, requiring less water for urine production (*17*). This mechanism aids water conservation in camels (*18*) and perhaps in water-deprived hibernators as well.

Our results provide the first direct evidence of a functional role for the gut microbiome in hibernation. UNS enables a symbiosis that provides TLGS and their gut microbes with the nitrogen required to build AAs and protein during the winter fast. Our demonstration of UNS as a mechanism for replenishing protein pools in TLGS has important implications beyond hibernation. For example, protein malnutrition *via* nitrogen-limited diets affects millions of people globally, a problem compounded by an increasing population and decreasing arable lands (*19*, *20*). Understanding the evolved mechanisms by which mammals maintain protein balance in nitrogen-limited situations may inform strategies for maximizing the health of other nitrogen-limited animals, including humans.

## Materials and Methods

### Experimental animals

This study used 47 13-lined ground squirrels (*Ictidomys tridecemlineatus*; TLGS). 39 of these (17 males, 22 females) were pups born in a vivarium on the University of Wisconsin-Madison (UW-Madison) campus to wild-caught female squirrels that were collected around Madison, Wisconsin in May 2018. These 39 TLGS were used in all analyses except for the urease activity assays (described below). The remaining 8 squirrels, used for urease activity assays, were wild-caught females.

Pregnant squirrels were housed individually at T_a_ of 22°C room with a 12:12 h light-dark cycle. Water and rodent chow (Teklad no. 2020X, Envigo, Indianapolis, IN, USA) were provided ad libitum, and diets were supplemented with fruit (apples) and sunflower seeds once per week. After parturition, pups remained with their mothers for 5 weeks and were then moved to individual cages. Following 2 weeks of ad libitum chow and fruit, food was restricted to 12 g chow per day (supplemented with 1 g of sunflower seeds once per week) to prevent excessive weight gain that can occur in squirrel pups born in captivity. The consistent diet among squirrels was important because diet can significantly impact the gut microbiome and the molar ratio of ^13^C to ^12^C in exhaled CO_2_. Squirrels were held under these conditions until the day of experiment (Summer TLGS group) or transferred to the cold room at the beginning of the hibernation season (Winter TLGS groups).

In mid-September, TLGS in the Winter groups were moved to a 4°C room in the UW-Madison Biotron, a controlled environment facility adjacent to the building that houses the research laboratory. The cold room was held in constant darkness except for ~5 min/day of dim light to enable activity checks. Food and water were removed after squirrels began regular torpor-arousal cycles, which was usually 24–72 h after being moved into the cold room.

The 39 squirrels were divided into three seasonal groups: Summer (14 squirrels), Early Winter (12 squirrels), and Late Winter (13 squirrels). Each seasonal group was divided into three treatment groups: ^13^C,^15^N-labeled urea with antibiotics, ^13^C,^15^N-labeled urea without antibiotics, and unlabeled urea without antibiotics. Mean days in hibernation for the Early Winter and Late Winter groups were 36 d (range 31–41 d) and 127 d (range 109–135 d), respectively. Mean body masses on experiment days for Summer, Early Winter, and Late Winter groups were 187.1 g (range 165.0–224.0 g), 155.6 g (range 123.4–187.2 g), and 134.1 g (range 105.5–167.0 g), respectively.

The UW-Madison Institutional Animal Care and Use Committee approved all procedures.

### Antibiotic treatment

To deplete gut microbes, antibiotic (ABX)-treated squirrels were administered a cocktail of ampicillin (1.0 g/L), vancomycin (0.5 g/L), neomycin (1.0 g/L) and metronidazole (1.0 g/L) in drinking water. Summer squirrels received ABX for 14 d prior to stable isotope breath analysis (see below). Squirrels in both Winter groups received ABX for 14 d prior to transfer to the cold room (4°C) in mid-September. To ensure the ABX remained effective until late winter, TLGS in the Late Winter group received ABX boosters once every six weeks for a total of three boosters per squirrel. On the day of ABX dosing, torpid TLGS were moved from the cold room to the lab (T_a_ 22°C). Approximately 6 h later when TLGS were euthermic (*22*), each was anesthetized (4% isoflurane-O_2_) and administered 1 ml of ABX water via oral gavage. After recovery from anesthesia, TLGS were moved back into the cold room. TLGS were artificially aroused for this procedure only if they were ≥75% through the length of their current torpor bout, which we estimated based on recent daily activity records. All TLGS successfully re-entered torpor within 24 h of this procedure.

### Urea treatment, live animal experiments, and stable isotope breath-testing

Each TLGS received two intraperitoneal (i.p.) injections of urea (^13^C,^15^N-labeled or unlabeled, 200 mg urea kg^−1^ body mass, 1 ml volume). TLGS were briefly anesthetized (4% isoflurane-O_2_ mixture) and urea was injected beneath the skin/abdominal muscle layers.

In Summer TLGS, the first urea injection occurred 7 days prior to breath analysis and the second occurred on the day of breath analysis, 3 h prior to tissue harvesting. Seven days was chosen for the first injection to allow sufficient time for urea nitrogen to be incorporated into the squirrels’ tissue protein pools and for ABX-treatment (given 14 days earlier) to deplete gut microbes in the ABX-treated subgroup. For TLGS in Early and Late Winter, instead of standardizing the time between first and second urea injection according to number of days, we chose to standardize according to approximate ‘metabolic time’. TLGS were artificially aroused from torpor for their initial urea injection, then placed back in the cold room to re-enter torpor. At some point within 14 days of re-entering torpor, the TLGS would arouse naturally (i.e., a natural interbout arousal, IBA) and then re-enter torpor. The TLGS would be artificially aroused 1 day following this re-entry into torpor and its breath analysis experiment would ensue as described for Summer TLGS (including the second urea injection). We chose this method to best standardize the time each TLGS spent at euthermic body temperature (T_b_), which we speculated was the metabolic state in which most microbial ureolytic activity and subsequent urea nitrogen incorporation would take place.

On the day of a breath analysis experiment, a TLGS was weighed and transferred to a Plexiglas holding box (25 × 10 × 10 cm in length, width, and height). For Summer TLGS, this transfer occurred once the animal arrived in the lab. For Early and Late Winter TLGS, this transferred occurred 3 h following their removal from the cold room, at which point they were fully euthermic (*22*). A Tygon tube connected the holding box to a cavity ring-down spectroscopy (CRDS) unit (Picarro, Sunnyvale, CA, USA) that measured the ^13^C:^12^C ratio (termed δ^13^C, expressed as ‰) in the squirrels’ exhaled CO_2_. Baseline breath measurements began once the TLGS was positioned in the box.

Air flow rate through the box was regulated by a vacuum pump to maintain a CO_2_ concentration of ~1%, the necessary concentration to make reliable δ^13^C measurements. Every 15 to 20 min, an air sample was automatically drawn into the CRDS unit for isotopic analysis. δ^13^C was calculated according to the equation:

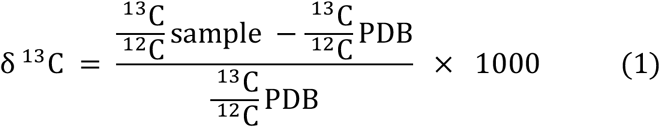

where PDB (Pee Dee Belemnite) served as a standard, and δ^13^C is expressed as parts per mil (‰).

Three baseline breath δ^13^C measurements were gathered over a ~45 min period. The TLGS was then anesthetized (4% isofluroane-O_2_ mixture), its rectal temperature taken, and the second urea i.p. injection administered. The TLGS was then returned to the CRDS box to recover and additional breath δ^13^C samples were gathered over the subsequent 3 h. At the 3 h mark, the TLGS was anesthetized, euthanized via decapitation, and its tissues were harvested.

### Tissue harvesting

Trunk blood was collected from each TLGS immediately following decapitation. Whole blood was collected in a 5 ml tube and centrifuged for 10 min at 10 000 g and 4°C to separate plasma and blood cell compartments. Plasma was collected and flash frozen in liquid N_2_.

Ceca were collected using sterilized dissection tools. Once isolated, ceca were weighed, sliced open, their contents collected in two aliquots (one for metabolomics analyses and one for metagenomic analyses), and both flash frozen in liquid N_2_. The cecum was rinsed in sterile phosphate-buffered saline to remove any remaining cecal content, excess liquid was gently scraped off, and the tissue cut into three pieces that were flash frozen and stored at −80°C. All steps involving ceca and cecal content were performed using sterile materials and equipment.

The entire liver and quadriceps muscle (right rear leg) were collected from each TLGS and flash frozen in liquid N_2_ for metabolomics analyses. Other tissues that were harvested but not included in the present study include heart, white adipose tissue, brown adipose tissue, kidney, and various intestinal segments and their contents including ileum, jejunum, proximal colon, and distal colon. All were flash frozen in liquid N_2_ for later analyses.

### Gut microbial urease activity

The eight wild-caught TLGS used in urease activity analyses were transferred to the cold room for hibernation in late September 2018. Late Winter TLGS (N=4) were sampled in February 2019. The Summer group (N=4) was sampled in April 2019, approximately two months after being moved to active season husbandry conditions.

Bacterial pellets from each TLGS cecal content sample were prepared according to (*23*). Pellets were flash frozen in liquid N_2_ and stored at −80°C.

On the day of analysis, bacterial pellets were thawed and mixed with 2-times volume assay buffer. Bacterial cells were lysed using sonication (3 × 10 s pulses at medium intensity; cooled on ice between pulses) and centrifuged for 30 min at 14 000 g and 4°C. The supernatant was collected and used immediately in a commercial urease activity assay kit (Sigma-Aldrich MAK120, St. Louis, MO, USA).

Urease activity was expressed as mg NH_3_ gram^−1^ content^−1^ h^−1^ to match the units reported in other urease studies.

### Cecal UT-B protein abundance

Cecal tissues for UT-B protein abundance analyses were harvested from captive-born TLGS used in a related project investigating gut microbial degradation of ^13^C-labeled carbohydrates *via* CRDS-based breath analysis. These TLGS were raised alongside and identical to the urea-treated TLGS, and the breath analysis procedure that preceded these animals’ tissue harvesting was identical to that of the urea-treated TLGS except that these TLGS were orally administered (via gavage) saline or saline-dissolved ^13^C-inulin (500 mg kg^−1^) or ^13^C-mannitol (150 mg kg^−1^). Importantly, though TLGS within treatment groups were administered different ^13^C-labeled substrates (Fig. S6A,B), we found no statistical evidence that substrate per se affected UT-B abundance. A two-way ANOVA for the Summer blot produced P-values of 0.750 and 0.008 for Substrate and Microbiome Presence, respectively (Fig. S6A; n=12); for the Late Winter blot, P-values were 0.110 and 0.003, respectively (Fig. S6B; n=12). With no significant effect of substrate on UT-B abundance, we proceeded to run immunoblots on Summer and Late Winter TLGS with intact microbiomes, irrespective of orally administered substrate (Fig. S6C). Moreover, use of these TLGS instead of urea-treated TLGS removed the potentially confounding effect of urea treatment on UT-B abundance.

Following the euthanasia and cecal harvesting procedures described previously, cecum samples were flash frozen and stored at −80°C. Cecal tissue lysates were prepared and then used for immunoblotting, where 30 μg of cecal tissue lysates were separated by SDS-PAGE, transferred to PVDF membrane, blocked with 5% nonfat milk, and incubated overnight with 1:1500 anti-UT-B antibody anti-SLC14A1 (Boster Biological Technology, Pleasanton, CA). Membranes were incubated with secondary antibody (1:2000) and immunoreactive bands were visualized by chemifluoresence and a UVP imaging system. Immunoblotting with anti-UT-B antibody produced a single protein band at approximately 48 kD (Fig. S6C; n=12), similar to the size reported for rat colon (*24*).

Using cecal lysates from microbiome-intact TLGS, specificity of the antibody for TLGS UT-B was confirmed by pre-incubation of antibody with the antigenic peptide A04926-BP (Boster Biological Technology, Pleasanton, CA) prior to immunoblotting, which eliminated the 48 kD band (Fig. S6D; n=10). Band intensity was quantified with ImageJ and values were normalized to intensity of individual Coomassie-stained sample lanes.

### Sample preparation for NMR

To determine whether urea-derived nitrogen and carbon are incorporated into host and microbial metabolomes, we performed ^13^C- and ^15^N-edited NMR-based metabolomics on cecal content, liver and quadriceps muscle.

For cecal content, ~50 mg of wet content was isolated, mixed with 8 homogenization beads and 350 μl of lysis buffer (10 mM phosphate buffer, pH 7.4), and homogenized using an Omni Bead Rupter 24 (Omni International, Kennesaw, GA, USA) through two 45 sec pulses (separated by 30 s) at 5.65 m s^−1^. The homogenized volume was then collected, and the beads were rinsed with an additional 50 μl of lysis buffer and vortexed for 10 s. This volume was added to the previous volume, bringing total volume up to ~400 μl. This volume was then centrifuged for 10 min at 10 000 *g* and 4°C. The supernatant (~300 μl) was collected and transferred to NMR tubes for NMR experiments (more details below).

For liver and quadriceps, ~50 mg of wet content was isolated, mixed with 8 homogenization beads and 600 μl of lysis buffer (10 mM phosphate buffer, pH 7.4), and the same homogenization, rinse, and centrifugation protocols described above were followed. After centrifugation, the recovered supernatant volume (usually ~550 μl) was diluted in 2-times volume ice-cold methanol to remove protein via precipitation, vortexed immediately for 30 s, and then incubated for 30 min at −20°C. Following the incubation period, samples were vortexed for 5 s and then centrifuged for 10 min at 10 000 *g* and 4°C. The supernatant was then collected and dried in a speed-vac for 40 to 45 h, while the protein pellet was frozen at −80°C for later protein quantification and isotope ratio mass spectrometry analyses. Following the speed-vac step, the dried sample was reconstituted in NMR buffer (details below) and transferred to NMR tubes for NMR experiments.

Different NMR buffers were used depending on the type of isotope-specific NMR analysis (^13^C or ^15^N) and the tissue being analyzed. The ^13^C-NMR buffer solvent was D_2_O, whereas the ^15^N-NMR buffer solvent was deionized H_2_O with 10% D_2_O added for locking the NMR spectrometer. Individual samples for ^13^C-NMR were adjusted to pH 7.4, whereas those for ^15^N-NMR were adjusted to pH 2.0. Finally, all buffers used included internal standards (0.25 mM trimethylsilyl propionate, 1 mM formate) and an antimicrobial agent (1 mM NaN_3_).

Within each tissue type, separate samples were prepared for ^13^C- and ^15^N analyses.

### NMR experiments and metabolomics analyses

All spectra were recorded on a Bruker Avance III spectrometer (Bruker, Billerica, MA, USA) operating at 600MHz (^1^H) and equipped with a cryogenically cooled triple resonance probe. The temperature of the samples was regulated to 278 K for experiments recorded on the samples in the ^15^N-NMR buffer and 298 K for experiments recorded on samples in the ^13^C-NMR buffer. NMR spectra on standard and experimental samples were recorded in an automated fashion using a Bruker sample jet with 3 mm NMR tubes and in-house written scripts. For each sample, optimal peak linewidths were achieved using an automated shimming protocol that starts with gradient shimming followed by a final optimization step directly on the DSS peak linewidth. In addition, the ^1^H 90degree pulse width was optimized for each sample. Once the automated optimization procedure was concluded, one-dimensional (1D) ^1^H spectra were recorded on each sample. In addition, for ^13^C analysis, ^13^C-edited 1D ^1^H spectra were recorded on the samples in the ^13^C-NMR buffer. For ^15^N analysis, two-dimensional (2D) ^1^H,^15^N-HSQC spectra were recorded on the samples in the ^15^N-NMR buffer.

For the 1D ^1^H spectra, suppression of the water signal was achieved using excitation sculpting (*25*) with pulsed field gradients and a perfect echo to improve the quality of the spectra (*26*). Each spectrum was recorded using a repetition delay of 3 seconds, a ^1^H spectral window of 16.0 ppm and 32,768 complex points for an acquisition time of 3.4 seconds. The ^1^H offset was centered on the water signal (4.76ppm). Each FID was accumulated with 128 scans for a total recording time of 14 minutes.

For ^13^C analysis, ^13^C-edited 1D ^1^H spectra were recorded using a pulse program that features an echo pulse train on proton (90°-delay-180°-delay-90°) with each delay set to 1/(2×^1^*J*_CH_), where ^1^*J*_CH_ is the one-bond coupling constant between ^1^H and ^13^C. Two broadband inversion pulses (BIP) (*27*) on ^13^C are given during the echo delays. Two experiments are then recorded in an interleaved manner. In the first experiment, the ^13^C BIP pulses are centered during each echo delay and magnetization from all protons is refocused. On the second experiment, the ^13^C BIP pulses are shifted such that, during the echo delays, the magnetization from protons attached to ^13^C evolves due to the ^1^*J*_CH_ coupling for a total period of 1/^1^*J*_CH_ and thus it is inverted at the end of the echo. On the other hand, protons attached to ^12^C are still unperturbed by these ^13^C BIP pulses. Subtraction of these two spectra will then result in cancellation of the signals from protons attached to ^12^C, leaving only signals from protons attached to ^13^C in the spectrum. This echo period is preceded by a weak presaturation pulse that is used to suppress signal from residual H_2_O and followed by a z-filter that cleans up any anti-phase magnetization prior to acquisition of the signals. ^13^C decoupling during ^1^H acquisition is achieved using bilevel decoupling (*28*) with CHIRP adiabatic pulses (*29*). These spectra were recorded using a repetition delay of 10 seconds, which includes the 5 second weak presaturation pulse, a ^1^H spectral window of 16.0 ppm and 4,096 complex points for an acquisition time of 0.426 seconds. The ^1^H offset was centered on the water signal (4.76ppm), while the ^13^C offset was set to 70ppm. Each FID was accumulated with 128 scans for a total time of 50 minutes required to record both interleaved spectra. These two interleaved spectra were saved in separate FIDs and subtracted using Bruker Topspin 3.5 software (Bruker Biospin).

For ^15^N analysis, 2D ^1^H,^15^N-HSQC spectra were recorded with a fast HSQC pulse program using a Watergate element for water suppression (*30*). The two hard 180degree pulses on ^15^N that are used in the pulse program were replaced by hyperbolic secant adiabatic inversion pulses with a 1ms duration and 190ppm bandwidth (*31*). Each spectrum was recorded using a repetition delay of 2 seconds. For ^15^N, the spectral window was set to 174.0 ppm, centered at 95 ppm, and 64 complex points were recorded. For ^1^H, the spectral window was set to 20.0 ppm, centered on the water signal at 4.76ppm, and 3,072 points were recorded, yielding an acquisition time of 0.246 seconds. ^15^N decoupling during ^1^H acquisition was achieved using a garp-16 decoupling sequence (*32*). Each FID was accumulated with 40 scans for a total recording time of 3 hours and 15 minutes for each HSQC spectrum.

All 1D ^1^H spectra were analyzed using the Chenomx software package (Version 8.5; Edmonton, AB, Canada). The 2D ^1^H,^15^N-HSQC spectra were processed with NMRPipe (*33*) and analyzed using NMRFAM-SPARKY software package (*34*). For ^15^N analysis, a set of 20 unlabeled amino acids at 10 mM concentration were used to prepare standard samples in the same ^15^N-NMR buffer at pH 2.0. 2D ^1^H,^15^N-HSQC spectra were recorded at natural abundance on each standard sample and used to identify the peaks in the tissue samples. Matching peaks were then integrated in NMRFAM-SPARKY.

### Isotope ratio mass spectrometry

To determine whether urea-derived nitrogen and carbon are incorporated into both host and microbial protein pools, we performed isotope ratio mass spectrometry (IRMS) on cecal content, liver, and quadriceps samples.

Protein precipitates were made for each sample as described in the NMR preparation process above. Each precipitate was dried in an 80°C oven for 24 h, and then ~3 mg of this dried precipitate was used for IRMS analysis.

Aliquots of dried samples (0.5 mg) for ^13^C and ^15^N analysis were placed in 3.5 × 5 mm tin cups (Costech, Valencia, CA). The tin cups were folded and placed in the Zero Blank auto-sampler inlet of an elemental analyzer (Costech, Valencia, CA, USA) and placed under helium. The specimens were dropped into a quartz furnace (1035°C) filled with chromium III oxide and silvered cobaltous/cobaltic oxide and a pulse of oxygen gas was introduced. The resulting gases were swept into a reduction furnace (copper wire, 650°C) in a stream of He and then directed into a G.C. separation column (65°C) using a ConFlo III, (Thermo Finnigan, Bremen, Germany). Effluent was directed through a quartz capillary and introduced into the ion source of a Delta V (Thermo Finnigan, Bremen, Germany) and the ion currents at m/z 28, 29, 30 followed by m/z 44, 45, and 46 employing a step change in the magnetic field strength. Aliquots of standard peach tree leaves (NIST Standard Material 1547, National Institute of Standards and Technology, US Department of Commerce, Gaithersburg, MD, USA) were analyzed before and after a batch of up to 40 bilk solids. The δ^13^C and δ^15^N.were calculated relative to the peach tree leaf standards. Precision for duplicates averaged 0.12 ‰ and 1.47 ‰ for δ^13^C and δ^15^N, where δ (‰) = (R_unknown_/R_standard_ −1)1000, and R equals the ratio of the heavy to light isotopes.

### Plasma/gut urea concentration

Urea concentrations in plasma and gut contents were determined using a spectrophotometric assay kit (Bioassay Systems, Hayward, CA, USA). Proximal colon contents (collected from adult female TLGS used for urease activity analysis) were centrifuged for 5 min at 400 g and 4°C and the supernatant was assayed according to kit instructions.

### Metagenomics

Genomic DNA from cecal content samples were extracted using the Qiagen DNeasy PowerSoil Kit (Qiagen, Hilden, Germany) according to manufacturer’s specifications with two modifications: (1) after adding C1 solution, tubes were put into a 65°C hot water bath for 10 min, and (2) before the C5 solution wash, samples were washed with 500 uL 100% ethanol. DNA was quantified using a Qubit Fluorometer (Invitrogen, San Diego, CA, USA). Eight samples (three summer, three early winter, and three late winter) were selected for community shotgun metagenomic sequencing based on individuals with the highest breath δ^13^C in each season. Sequencing was conducted on an Illumina NovaSeq6000 (Illumina, San Diego, CA, USA) using an S4 Reagent Kit v1.5 at 2×150bp PE. Fastq files were submitted to the NCBI sequence read archive under BioProject PRJNA693524.

For quality control, raw reads were run through Trimmomatic v0.38 (*35*) to remove sequencing adapters and low-quality reads. Host DNA sequences were removed using bowtie2 v2.2.2 (*36*) by aligning all reads against the TLGS genome (GenBank and RefSeq assembly accession = GCA_000236235.1). Reads that aligned to the host genome were removed using samtools v0.1.19 (*37*, *38*). FastQC v0.11.8 (*39*) was used after each step to evaluate read quality.

Reads were then assembled using metaSPAdes v3.13.0 (*40*) and assembly statistics calculated using MetaQUAST v5.0.2 (*41*) and bowtie2 (*36*). Open reading frames (ORFs) were predicted using prodigal v2.6.3 (*42*). Code for quality control through open reading frame prediction can be found at https://github.com/ednachiang/Urea.

Translated proteins from the ORF predictions were then compared against the Pfam profiles for proteins (*43*) involved in urea recycling (urease structural and accessory proteins, see Fig. 4C) as defined by KEGG (*44*) using HMMER (*45*). Identified sequences were then mapped back to their contig of origin and taxonomically classified using centrifuge (*46*).

### Area under the curve (AUC) calculations for breath-testing experiments

To calculate the total breath δ^13^C values over 3 h (Fig. 3D), we used trapezoidal numerical integration with 15 min intervals according to the previously reported method (*47*). The δ^13^C values were first rescaled so that zero corresponded to the most negative δ^13^C value. The values for time were rescaled so that zero corresponded to the first time point, defined as t=0 at i=1. The average δ^13^C values at any given time point for all individuals in a given treatment were used as the data point for each corresponding time point of measurement. To compute the area under the curve, the midpoint method was used for numerical integration with variable time intervals as the equation:

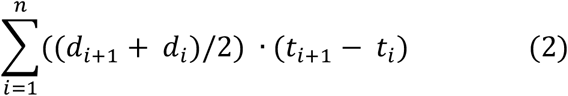

where d is the δ^13^C value and t is time.

### Statistical analyses

Plasma urea concentrations were compared using 1-way ANOVA with Tukey multiple comparisons test (Fig. 2A). UT-B protein expression was compared using 2-way ANOVA with Tukey multiple comparisons test (Fig. 2B). Seasonal AUC breath δ^13^C values for TLGS with intact microbiomes were compared using 1-way ANVOA with Tukey multiple comparisons test, while seasonal differences in breath AUC for TLGS with intact or depleted microbiomes were analyzed using unpaired t-tests (Fig. 3D). Urease gene percentages in TLGS with intact microbiomes were compared using 1-way ANOVA with Tukey multiple comparisons test (Fig. 3E). Seasonal ^15^N-Protein, -Metabolome and -Compound values for TLGS with intact microbiomes were compared using 1-way ANVOA with Tukey multiple comparisons test, while within seasons, mean ^15^N-Protein, -Metabolome and -Compound values for TLGS with intact or depleted microbiomes were compared using unpaired t-tests (Fig. 4). For data presented in whisker plots, the upper and lower bounds of the box represent 75^th^ and 25^th^ percentiles, respectively; the horizontal line within the box represents the median value, and the upper and lower whiskers represent the maximum and minimum values within the data set, respectively. Statistical significance was set at P<0.05 for all comparisons, and GraphPad Prism 9.0 (GraphPad Software, San Diego, CA, USA) was used to run all comparison and create all graphs.

## Acknowledgments

We thank Mike Grahn, Adam Steinberg, Samantha Cailey and Profs. Warren Porter and Sandra Martin for assistance with aspects of this study.

## Author contributions

HVC, FMAP and GS conceived the study; HVC, FMAP, GS, MDR and MT developed the methods; MDR, EC, MT, SG and KV collected data; MDR, FMAP, GS, EC and YL conducted analyses; MDR wrote manuscript with input from all authors.

## Data and materials availability

Metabolomics data available from corresponding author upon request. Metagenomic data available at NCBI, Bioproject PRJNA693524.

## Supplementary Materials

Materials and Methods, Figs. S1-S7, Tables S1-S3, References (22-47).

