## Supplementary materials for "Urea nitrogen recycling via gut symbionts increases in hibernators over the winter fast"

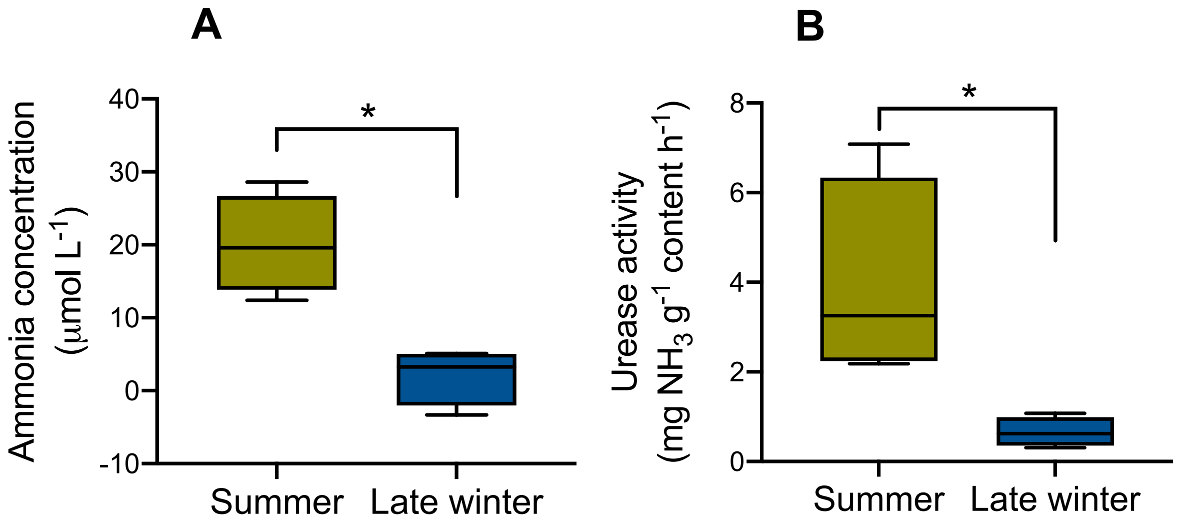


**Fig. S1. Gut microbial ammonia concentrations and urease activity levels.** Mean (A) [ammonia] and (B) urease activity levels in bacterial pellets isolated from cecal contents of non-urea-treated TLGS (t-test, P=0.030 (A), P=0.004 (B); n=4 for each seasonal group).

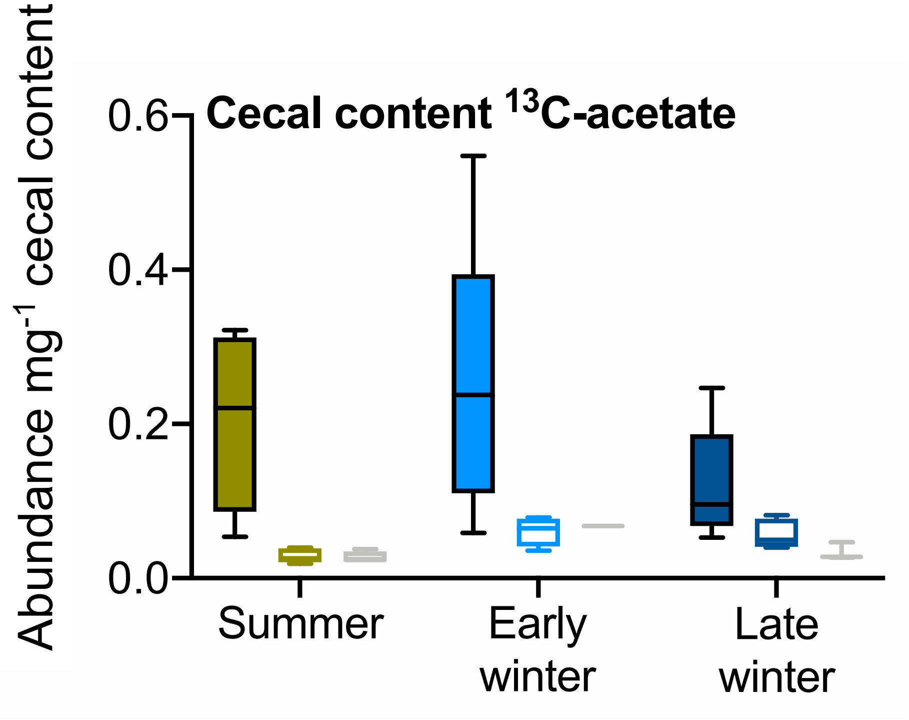


**Fig. S2. $^{13}$C-acetate abundance in cecal content.** Mean values are arbitrary units expressed per milligram of cecal content in TLGS treated with $^{13}$C,$^{15}$N-urea (grey bars for TLGS treated with unlabeled urea). Solid bars for TLGS with intact gut microbiomes, open bars for TLGS with depleted microbiomes. Data for $^{13}$C,$^{15}$N-urea-treated TLGS compared with 2-way ANVOA (Season P=0.443; Microbiome P=0.0002; Interaction P=0.534; n=4-5 for labeled urea groups, n=2-4 for unlabeled urea groups).

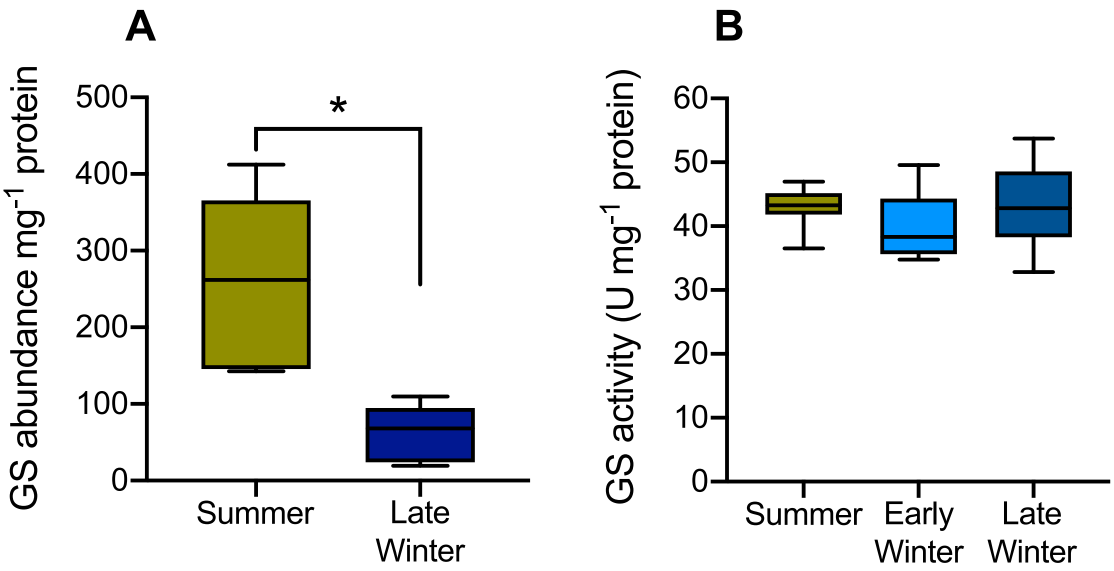


**Fig. S3. Liver glutamine synthetase (GS) abundance and activity.** (A) GS abundance per mg protein and (B) activity levels in livers of microbiome-intact TLGS. Asterisk in (A) indicates significant difference between groups (t-test, P=0.007). Values in (B) compared using 1-way ANOVA (P=0.258). n=7 for all seasonal groups in (A) and (B). TLGS used in analyses had intact gut microbiomes and were treated with labeled or unlabeled urea.

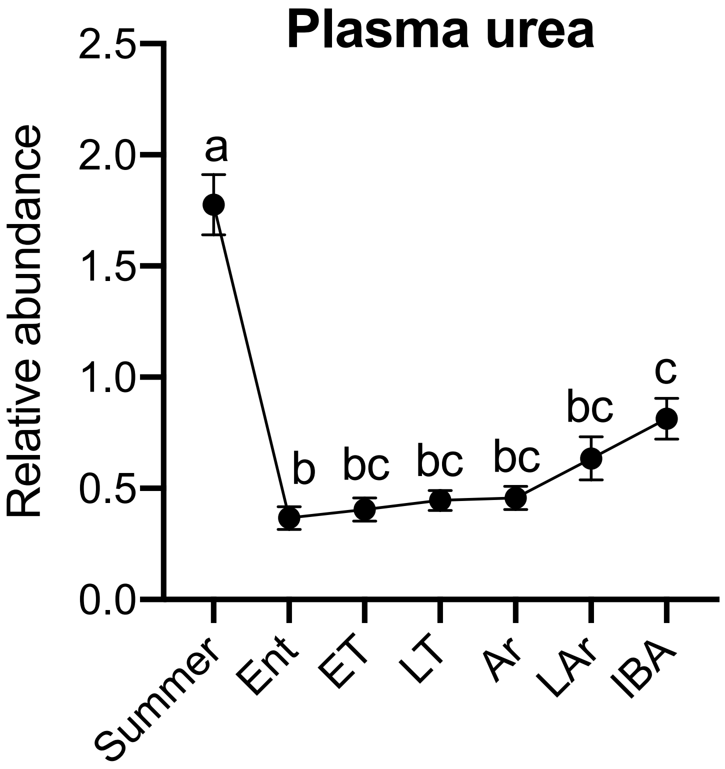


**Fig. S4. Mean relative abundance of urea in plasma of TLGS at different stages of the torpor-arousal cycle.** Blood plasma from animals sampled during summer active season (Summer), entry into torpor (Ent), early torpor (ET), late torpor (LT), early arousal (Ar), late arousal (LAr) and middle of interbout arousal (IBA). Values compared using 1-way ANVOA with Tukey multiple comparisons test ($P<0.0001$, n=6-8). Values that do not share a letter are significantly different ($P<0.05$). Data from Epperson LE, Karimpour-Fard A, Hunter LE, Martin SL. Metabolic cycles in a circannual hibernator. *Physiological genomics*. 2011 Jul;43(13):799-807. See that paper for details.

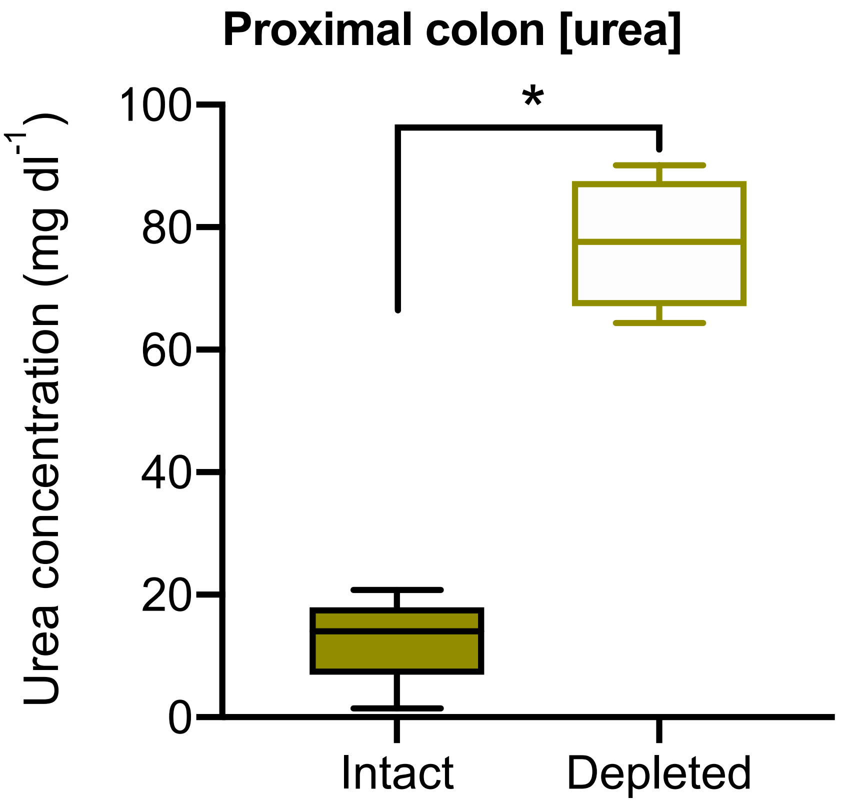


**Fig. S5**. **Urea concentrations in proximal colon of TLGS with intact and depleted gut microbiomes.** Data are from TLGS treated with $^{13}C$,$^{15}N$-urea in the summer active season (t-test, P<0.0001, n=4-5).

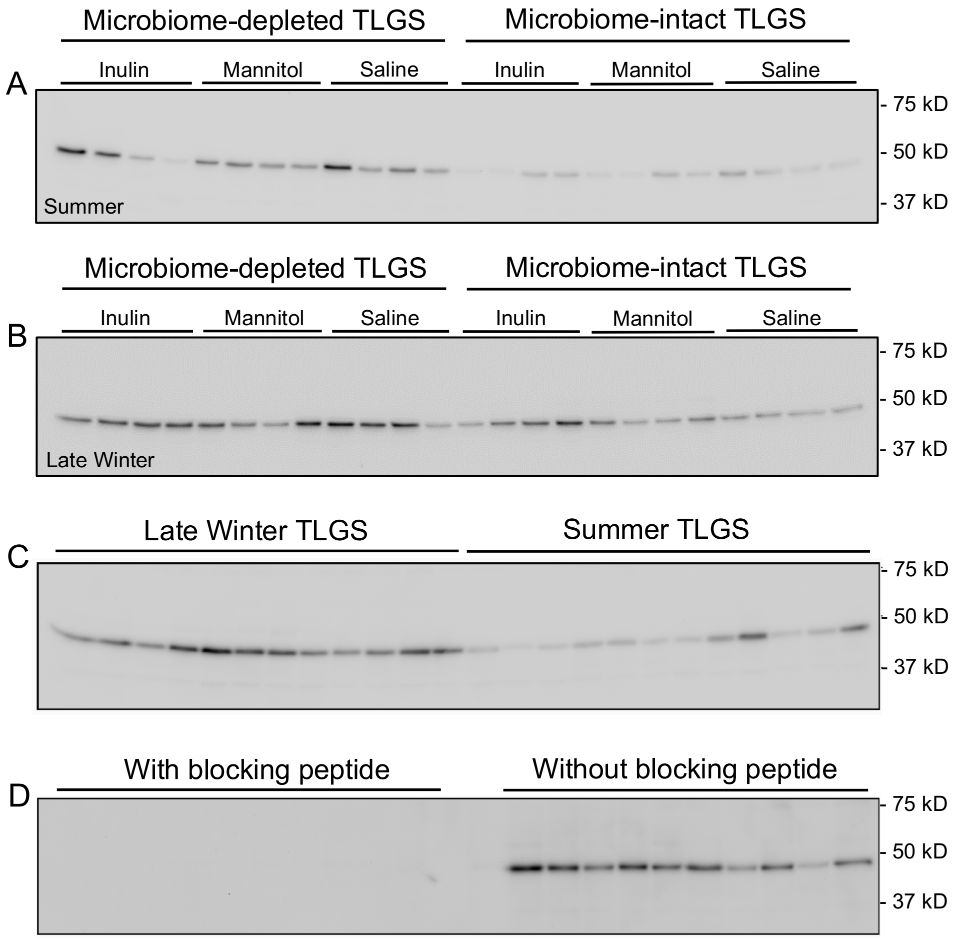


**Fig. S6. Urea transporter (UT-B) immunoblots for TLGS cecal tissue lysates**. Lysates for (A) Summer and (B) Late Winter TLGS were made from TLGS used in a related study that involved oral treatment of the substrates listed atop (A) and (B) images. Substrate had no effect on Coomassie-normalized UT-B abundance, but gut microbiome presence did ((A) Substrate factor P=0.75, Microbiome factor P=0.008, Interaction P=0.556; (B) Substrate factor P=0.11, Microbiome factor P=0.003, Interaction P=0.877). Data for (A) and (B) are plotted and compared in Fig. 2B. (C) UT-B immunoblots for cecal lysates from Summer and Late Winter TLGS with intact microbiomes, (D) Antibody specificity confirmation experiments completed with and without antigenic blocking peptide using cecal lysates from TLGS with intact gut microbiomes.

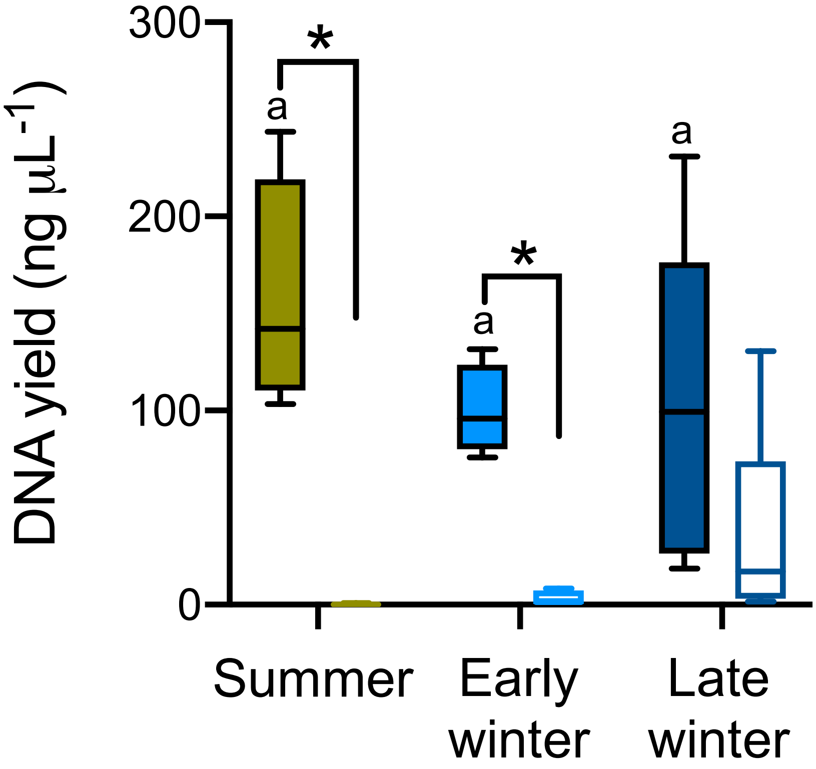


**Fig. S7. DNA extraction yields from microbiome-intact (solid bars) and -depleted (open bars) TLGS across seasons.** Asterisks indicate a significant difference (t-test, $P < 0.05$) between microbiome-intact and -depleted samples within a season. The same letter indicates no significant difference between microbiome-intact seasonal groups. n=5 for each group except Early Winter microbiome-depleted group, where n=4.

**Table S1.** P-values for statistical tests shown in Fig. 4 for $^{15}$N-enriched compounds in cecum content, liver and muscle of 13-lined ground (TLGS) treated with $^{13}$C,$^{15}$N-labeled urea (see text for details). Bolded values represent statistically significant findings for respective test (P<0.05).

| | | **2-way ANOVA** | | | **1-way ANOVA** | | | **t-test** | | |
|---|---|---|---|---|---|---|---|---|---|---|
| **Tissue** | **Feature** | **Interact.** | **Season** | **GM** | **S-EW** | **S-LW** | **EW-LW** | **S** | **EW** | **LW** |
| Cecum content | 15N-Metabolome | **0.0024** | 0.3604 | **<0.0001** | **0.0466** | **0.0348** | 0.9853 | **0.0002** | **0.0027** | **0.0311** |
| | 15N-Ammonia | **0.0347** | 0.3303 | **<0.0001** | 0.1405 | 0.1413 | 1.0000 | **0.0030** | **0.0008** | 0.1902 |
| | 15N-Glutamine | 0.1479 | 0.177 | **<0.0001** | 0.4681 | 0.1207 | 0.6272 | **0.0001** | **0.0029** | **0.0006** |
| | 15N-Alanine | 0.8976 | 0.689 | **<0.0001** | 0.9268 | 0.9837 | 0.8557 | **0.0006** | **0.0070** | **0.0164** |
| | 15N-Lysine | **0.0031** | 0.436 | 0.0619 | 0.2516 | **0.0132** | 0.2347 | **0.0003** | 0.7405 | 0.4305 |
| | 15N-Valine | 0.2638 | 0.6041 | 0.1367 | 0.3268 | 0.0890 | 0.6878 | **0.**5186 | 0.1367 | 0.2266 |
| Liver | 15N-Protein | **0.0108** | **<0.0001** | **<0.0001** | **0.0206** | **0.0001** | **0.0233** | **0.0015** | **0.0007** | **0.0014** |
| | 15N-Metabolome | **0.0049** | **<0.0001** | **<0.0001** | **0.0014** | **0.0001** | 0.2053 | 0.8873 | **0.0114** | **0.0002** |
| | 15N-Ammonia | **0.0018** | **<0.0001** | **<0.0001** | **0.0045** | **0.0000** | **0.0024** | 0.1134 | 0.1611 | **0.0001** |
| | 15N-Glutamine | 0.2348 | **0.0086** | **0.0002** | 0.2722 | **0.0232** | 0.3378 | 0.0779 | **0.0144** | **0.0228** |
| | 15N-Alanine | **0.0268** | 0.0559 | **0.0002** | 0.2642 | **0.0264** | 0.3820 | 0.4991 | 0.0729 | **0.0013** |
| | 15N-Lysine | 0.1689 | **0.0281** | 0.4774 | 0.1727 | **0.0059** | 0.1724 | 0.4768 | 0.8735 | 0.0700 |
| Muscle | 15N-Protein | 0.0837 | **0.0003** | **<0.0001** | 0.0970 | **0.0009** | **0.0475** | 0.1178 | **0.0006** | **0.0185** |
| (quadriceps) | 15N-Metabolome | 0.521 | 0.5368 | **0.0155** | 0.9725 | 0.4140 | 0.5379 | 0.2930 | 0.0282 | 0.2605 |
| | 15N-Ammonia | 0.6239 | 0.4201 | **0.0494** | 0.6432 | 0.3818 | 0.8879 | 0.6204 | 0.0718 | 0.2650 |
| | 15N-Glutamine | 0.9657 | 0.5265 | 0.1594 | 0.9580 | 0.6198 | 0.7604 | 0.5276 | 0.5150 | 0.2485 |
| | 15N-Alanine | 0.8463 | 0.6559 | 0.2669 | 0.9150 | 0.9656 | 0.9876 | 0.6033 | 0.3686 | 0.7188 |
| | 15N-Lysine | 0.3715 | 0.7438 | 0.4774 | 0.9967 | 0.6425 | 0.6890 | 0.3971 | 0.9844 | 0.2122 |

*2-way ANOVA: Comparison of $^{15}$N-compound abundances across two factors: season (Summer, Early Winter, Late Winter) and gut microbiome (GM; intact, depleted). Values represent effect of season, gut microbiome (GM) and interaction. 1-way ANOVA: Used to determine the effect of season on 15N-compound abundance in microbiome-intact TLGS tissues. Values represent Tukey multiple comparisons results for comparisons between Summer (S), Early Winter (EW) and Late Winter (LW). t-test: Used to determine the effect of a gut microbiome on $^{15}$N-compound abundance in TLGS from Summer, Early Winter and Late Winter. Values are P-values when comparing TLGS with intact and depleted gut microbiomes within a season. $^{15}$N-Protein portrays $^{15}$N-incorporation into protein content as determined by isotope ratio mass spectrometry. $^{15}$N-Metabolome portrays $^{15}$N-incorporation into the total pool of compounds, both identifiable and non-identifiable, using $^{15}$N-edited NMR.*

**Table S2.** Data for individual 13-lined ground squirrels (TLGS) from control group, meaning these animals had intact gut microbiomes and were treated with unlabeled urea. Features are the same as those defined in Table S1 caption except for Breath $\delta^{13}C$ AUC, which represents the area under the curve calculated from each animal's breath $\delta^{13}C$ vs. time graph. Breath $\delta^{13}C$ AUC values can be compared with those for other treatment groups shown in Fig. 3D, and all other features can be compared with those for other treatment groups shown in Fig. 4.

| | | **Summer** | | | | **Early Winter** | | **Late Winter** | | |
|---|---|---|---|---|---|---|---|---|---|---|
| **Tissue** | **Feature** | S64 | S65 | S67 | S70 | EW8 | EW12 | W58 | W59 | W61 |
| Breath | $\delta$13C AUC | -51.35 | -61.01 | -50.11 | -56.89 | -67.01 | -63.27 | -68.64 | -62.66 | -68.02 |
| Cecum content | 15N-Protein | 3.84* | | | | 9.56* | | | | |
| | 15N-Ammonia | 0.76 | 0.96 | 0.96 | 1.69 | 1.59 | 0.67 | 1.07 | 0.94 | 0.54 |
| | 15N-Glutamine | 0.20 | 0.20 | 0.61 | N/A | 0.49 | 0.58 | 0.51 | 0.34 | 0.11 |
| | 15N-Alanine | N/A | N/A | N/A | N/A | N/A | N/A | 1.27 | 0.15 | N/A |
| | 15N-Lysine | N/A | N/A | 0.11 | 0.23 | 0.53 | N/A | 0.03 | 0.72 | N/A |
| | 15N-Valine | N/A | N/A | N/A | 0.08 | N/A | N/A | N/A | N/A | N/A |
| Liver | 15N-Protein | 6.77 | 6.71 | 6.80 | 6.83 | 6.90 | 7.49 | 7.71 | 7.33 | 7.25 |
| | 15N-Ammonia | 0.22 | 0.27 | 0.23 | 0.27 | 0.32 | 0.21 | 0.61 | 0.39 | 0.29 |
| | 15N-Glutamine | 0.16 | 0.27 | N/A | N/A | 0.76 | 0.65 | 0.70 | 0.52 | 0.64 |
| | 15N-Alanine | 0.30 | 0.36 | 0.48 | 0.30 | 0.25 | 0.22 | 0.47 | 0.32 | 0.29 |
| | 15N-Lysine | N/A | N/A | 0.14 | 0.62 | 0.47 | 0.61 | 0.95 | 0.77 | 0.67 |
| Muscle | 15N-Protein | 5.15 | 5.07 | 5.13 | 5.29 | 5.85 | 6.03 | 6.18 | 5.87 | 5.86 |
| | 15N-Ammonia | 0.23 | 0.28 | 0.17 | 0.23 | 0.31 | 0.14 | 0.21 | 0.19 | 0.50 |
| | 15N-Glutamine | 0.37 | N/A | 0.26 | 0.33 | 0.44 | 0.52 | N/A | 0.15 | 0.43 |
| | 15N-Alanine | 0.50 | 0.34 | 0.24 | 0.38 | 0.29 | 0.09 | 0.34 | 0.18 | 0.30 |
| | 15N-Lysine | 0.75 | 0.96 | 0.75 | 0.90 | 1.22 | 0.72 | 0.68 | 1.62 | 1.00 |

*Values with asterisks for Cecum content–$^{15}N$-Protein represent measurements made on pooled samples from each TLGS in the treatment group. Protein pellet for Late Winter cecum content was insufficiently sized for analysis. Values labeled as N/A represent measurements that were below the detection limit of the $^{1}H$-$^{15}N$ spectroscopy method and thus effectively absent from the sample.*

**Table S3.** Metagenome summary statistics. The first letter in sample names represents the season in which the individual was sampled: S = Summer, E = Early Winter, W = Late Winter.

| Sample | Raw sequences | Sequences after quality control | Assembly number of contigs | Assembly number of contigs >= 1000bp | Assembly total length (bp) | Assembly total length (contigs >= 1000bp) | Assembly N50 (bp) | Assembly L50 (bp) | Input reads alignment rate to assembly (%) |
|---|---|---|---|---|---|---|---|---|---|
| S63 | 72,780,964 | 64,434,082 | 850,372 | 93,487 | 653,702,712 | 353,599,221 | 1231 | 69,287 | 91.83 |
| S66 | 77,365,266 | 69,988,682 | 953,255 | 116,375 | 761,383,630 | 424,297,547 | 1319 | 78,189 | 92.06 |
| S80 | 48,874,842 | 44,667,712 | 606,266 | 65,120 | 464,757,973 | 250,421,984 | 1234 | 48,813 | 91.38 |
| E02 | 49,017,170 | 45,972,818 | 562,935 | 67,861 | 474,963,737 | 272,305,707 | 1452 | 38,608 | 92.65 |
| E06 | 79,323,828 | 64,434,082 | 580,011 | 80,990 | 488,658,189 | 282,668,497 | 1412 | 48,443 | 95.43 |
| E07 | 111,027,340 | 102,160,254 | 436,174 | 67,638 | 470,301,924 | 321,277,065 | 3238 | 15,636 | 97.10 |
| W56 | 56,408,336 | 31,110,400 | 403,747 | 48,077 | 310,570,829 | 167,532,819 | 1198 | 36,857 | 89.93 |
| W62 | 56,036,528 | 49,705,466 | 280,112 | 37,760 | 292,138,335 | 193,729,750 | 3343 | 9,097 | 96.28 |
| W64 | 63,680,326 | 47,916,308 | 392,526 | 44,172 | 338,339,800 | 198,822,874 | 1746 | 21,082 | 93.32 |
